# Gray matter covariations and core symptoms of autism. The EU-AIMS Longitudinal European Autism Project

**DOI:** 10.1101/2020.06.26.171827

**Authors:** Ting Mei, Alberto Llera, Dorothea L. Floris, Natalie J. Forde, Julian Tillmann, Sarah Durston, Carolin Moessnang, Tobias Banaschewski, Rosemary J. Holt, Simon Baron-Cohen, Annika Rausch, Eva Loth, Flavio Dell’Acqua, Tony Charman, Declan G. M. Murphy, Christine Ecker, Christian F. Beckmann, Jan K. Buitelaar, the EU-AIMS LEAP group

## Abstract

**Background:** Voxel-based Morphometry (VBM) studies in Autism Spectrum Disorder (autism) have yielded diverging results. This might partly be attributed to structural alterations being associating with the combined influence of several regions rather than with a single region. Further, these structural covariation differences may relate to continuous measures of autism rather than with categorical case-control contrasts. The current study aimed to identify structural covariation alterations in autism, and assessed canonical correlations between brain covariation patterns and core autism symptoms.

**Methods:** We studied 347 individuals with autism and 252 typically developing individuals, aged between 6 and 30 years, who have been deeply phenotyped in the Longitudinal European Autism Project (LEAP). All participants’ VBM maps were decomposed into spatially independent components using Independent Component Analysis. A Generalized Linear Model (GLM) was used to examine case-control differences. Next, Canonical Correlation Analysis (CCA) was performed to separately explore the integrated effects between all the brain sources of gray matter variation and two sets of core autism symptoms.

**Results:** GLM analyses showed significant case-control differences for two independent components. The first component was primarily associated with decreased density of bilateral insula, inferior frontal gyrus, orbitofrontal cortex, and increased density of caudate nucleus in the autism group relative to typically developing individuals. The second component was related to decreased densities of the bilateral amygdala, hippocampus, and parahippocampal gyrus in the autism group relative to typically developing individuals. The CCA results showed significant correlations between components that involved variation of thalamus, putamen, precentral gyrus, frontal, parietal, and occipital lobes, and the cerebellum, and repetitive, rigid and stereotyped behaviors and abnormal sensory behaviors in autism individuals.

**Limitations:** Only 55.9% of the participants with autism had complete questionnaire data on continuous parent-reported symptom measures.

**Conclusions:** Covaried areas associated with autism diagnosis and/or symptoms are scattered across the whole brain and include the limbic system, basal ganglia, thalamus, cerebellum, precentral gyrus, and parts of the frontal, parietal, and occipital lobes. Some of these areas potentially subserve social-communicative behavior whereas others may underpin sensory processing and integration, and motor behavior.

## Background

Autism spectrum disorder (autism) is an early onset neurodevelopmental condition characterized by core deficits in social interaction and communication, along with restrictive interests and behavior, and sensory abnormalities (1). Magnetic resonance imaging (MRI) studies have increased our understanding of the neuroanatomical underpinnings of autism and show that autism is associated, at the group level, with brain structural changes (2). However, many results are not robust across different studies. For example, two studies using the same large-scale open access Autism Brain Imaging Data Exchange (ABIDE) dataset (3) came to different conclusions with regard to the volume of the pallidum (4, 5). Also, across whole brain approaches investigating cortical (i.e. cortical thickness and surface area) and subcortical (i.e. volume) features have been inconsistent; two large-scale pooled estimate analytical studies observed diverging results of cortical changes in autism (6, 7). Similarly, autism studies quantifying voxel-wise gray matter (GM) density also found divergent results of GM differences between autism diagnosed and control individuals; for instance, meta-analyses reported diverse changes of GM morphometry in autistic individuals on average, reporting either increased or decreased density of right inferior temporal gyrus in autism (8, 9). Even when taking age into account, studies still observed different structural brain alterations in children and adolescents with autism (10, 11).

A commonality to all these studies is their reliance on mass-univariate statistics. This approach identifies alterations in isolated regions or voxels but ignores possible relationships between them. The brain is a complex system of interconnected networks, and research into the neural basis of autism has moved away from focusing on local abnormalities into conceptualizing autism as a disorder of alterations in structural and functional brain connectivity (12). This implies that structural brain alterations in autism likely reflect the combined influence of several regions and are not confined to one specific region (13, 14). The present paper aims to advance prior work on brain structural neural correlates of autism in two ways. First, we aim to move away from the standard univariate approach and incorporate an alternative that adheres more closely to the hypothesis of autism as a disconnection syndrome (13), thus providing greater sensitivity for between-group effects. For this purpose, we identify inter-regional sources of structural covariation using Independent Component Analysis (ICA) (15), a data-driven unsupervised approach that allows the identification of interconnected brain regions across the whole brain. It has previously been applied successfully to identify covariance of brain morphometry in patients with psychiatric disorders (16–18). Second, we move beyond the categorical autism case-control comparison towards exploring associations between brain structure and symptom dimensions or profiles of autism. Although former studies have used univariate approaches to explore the relationship between brain substrates and clinical phenotypes (6, 19), such associations are potentially the consequence of integrated effects across multiple symptoms dimensions and brain regions, rather than simple associations between a specific brain region and a specific symptom dimension. To study such multidimensional associations multivariate methods are effective (20, 21) and here we achieve this integration using Canonical Correlation Analysis (CCA) (22).

In summary, we investigate alterations in GM morphometric covariations in a deeply phenotyped large European autism case-control sample (23, 24) that allows us to improve our understanding of correlated structural brain alterations in autism. Subsequently, we focus on the covariation between the identified structural features and symptom behavior profiles among individuals with autism.

## Methods

### Participants

The participants were selected from the first wave of the European Autism Interventions—A Multicentre Study for Developing New Medications (EU-AIMS) Longitudinal European Autism Project (LEAP) dataset, which is a large multicenter study that aims to identify and validate biomarkers for autism (24). In total, six centers are involved: Institute of Psychiatry, Psychology and Neuroscience, King’s College London, United Kingdom; Autism Research Centre, University of Cambridge, United Kingdom; Radboud University Medical Centre, Nijmegen, the Netherlands; University Medical Centre Utrecht, the Netherlands; Central Institute of Mental Health, Mannheim, Germany; and University Campus Bio-Medico, Rome, Italy. Each participant underwent clinical, cognitive, and MRI assessment. Autism diagnoses were confirmed by clinicians according to the Diagnostic and Statistical Manual-IV (DSM-IV), International Statistical Classification of Diseases and Related Health Problems 10th Revision (ICD-10), or DSM-5. The study was approved by local ethical committees in each participating center, and written informed consent was provided by all participants and/or their legal guardians (for those<18 years old). For further details on experimental design and clinical assessments, see (23, 24).

In the present study, we selected participants with available structural MRI data. All images were inspected visually and subjects were excluded in cases of brain injury or structural abnormalities (e.g. enlarged ventricles or cysts), excessive head motion, or preprocessing failure (n=29). We excluded the participants from the Rome site due to low sample size (n=1). We also excluded participants without full-scale intelligence quotient (FSIQ) data in the further statistical analyses (n=5). This resulted in a sample of 599 participants from 5 sites, including 347 individuals with autism and 252 typically developing (TD) controls. Demographic and clinical information is shown in Table 1.

**Table 1.**
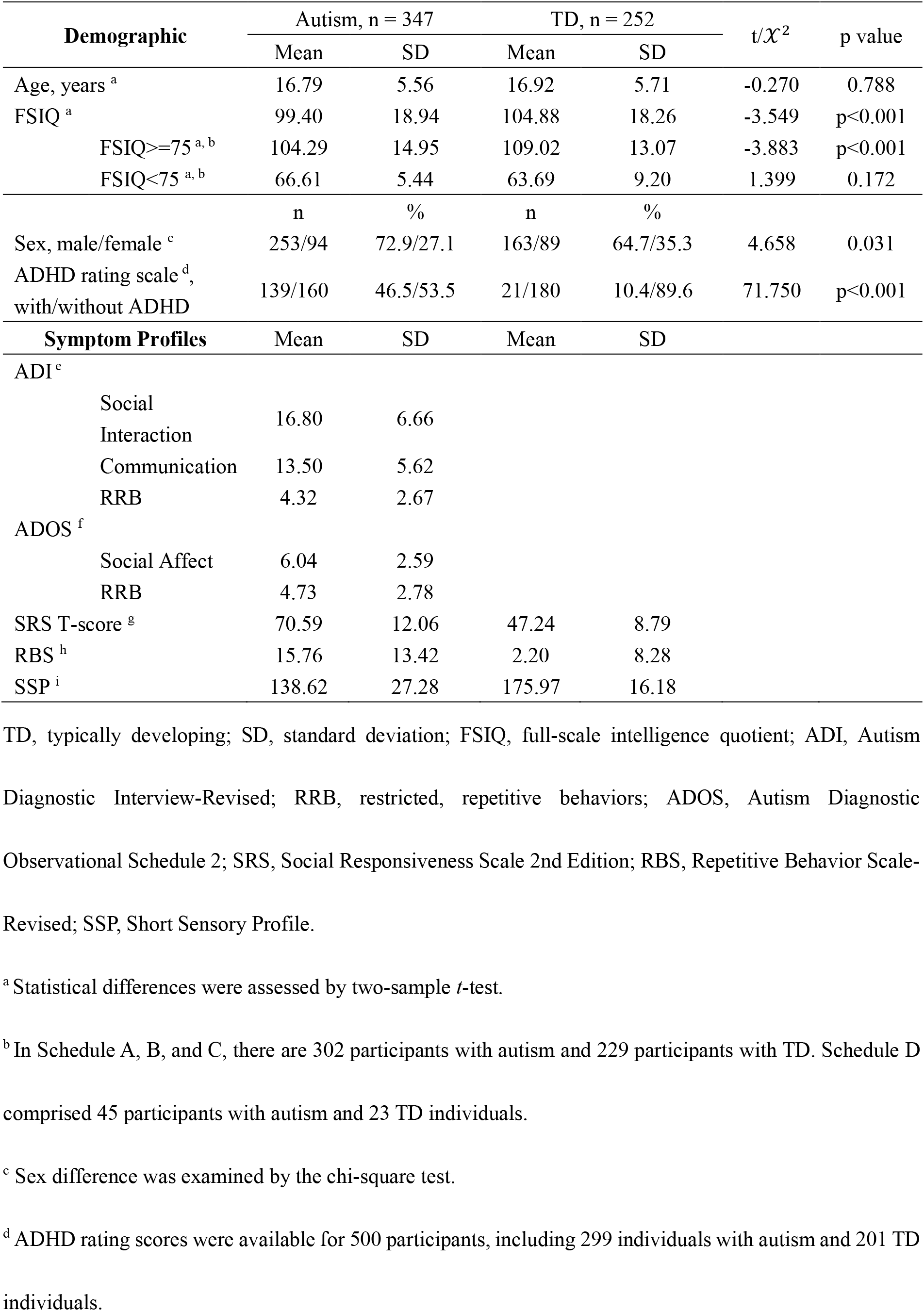
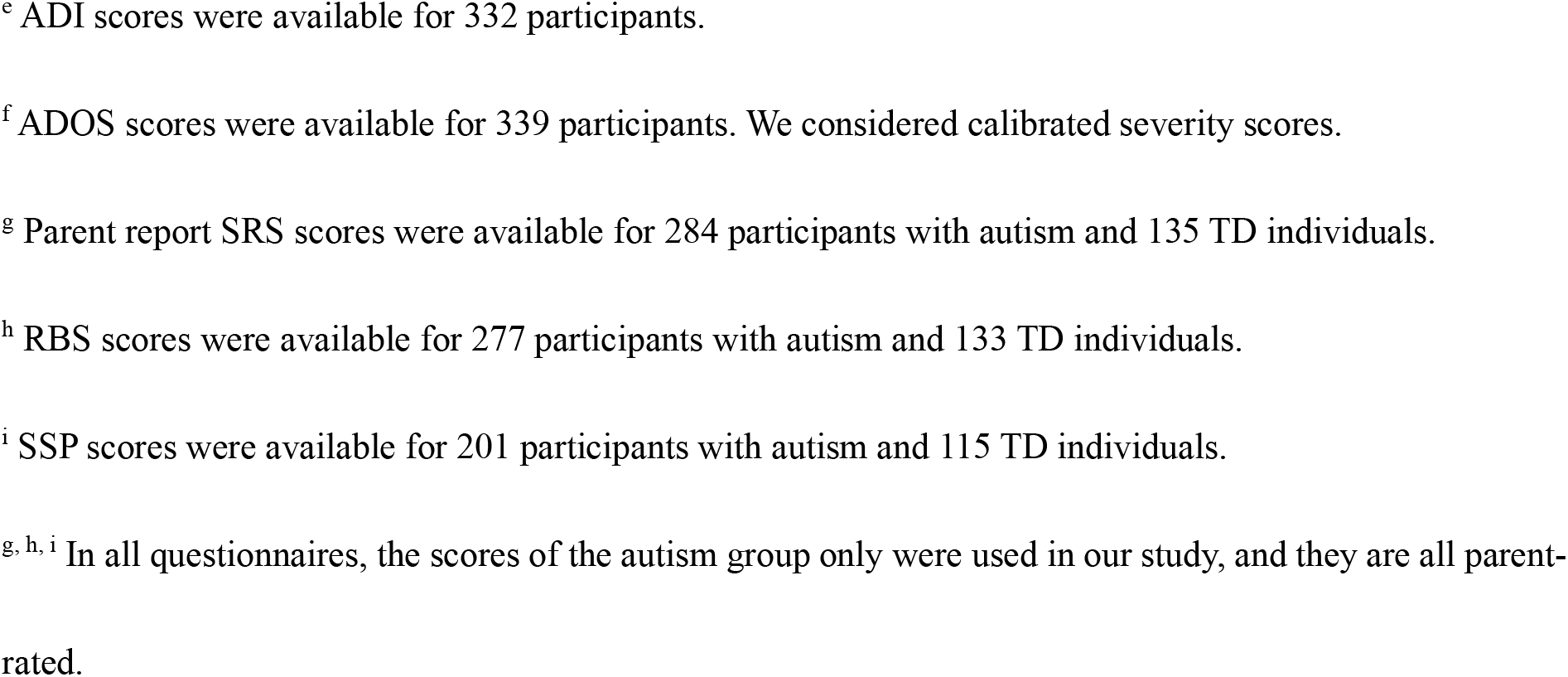
Participant characteristics

| Demographic | Autism, n = 347 | | TD, n = 252 | | t/$\chi^2$ | p value |
| --- | --- | --- | --- | --- | --- | --- |
|  | Mean | SD | Mean | SD |  |  |
| Age, years <sup>a</sup> | 16.79 | 5.56 | 16.92 | 5.71 | -0.270 | 0.788 |
| FSIQ <sup>a</sup> | 99.40 | 18.94 | 104.88 | 18.26 | -3.549 | p<0.001 |
| FSIQ≥75 <sup>a, b</sup> | 104.29 | 14.95 | 109.02 | 13.07 | -3.883 | p<0.001 |
| FSIQ<75 <sup>a, b</sup> | 66.61 | 5.44 | 63.69 | 9.20 | 1.399 | 0.172 |
|  | n | % | n | % |  |  |
| Sex, male/female <sup>c</sup> | 253/94 | 72.9/27.1 | 163/89 | 64.7/35.3 | 4.658 | 0.031 |
| ADHD rating scale <sup>d</sup>,<br>with/without ADHD | 139/160 | 46.5/53.5 | 21/180 | 10.4/89.6 | 71.750 | p<0.001 |
| Symptom Profiles | Mean | SD | Mean | SD |  |  |
| ADI <sup>e</sup> |  |  |  |  |  |  |
| Social Interaction | 16.80 | 6.66 |  |  |  |  |
| Communication | 13.50 | 5.62 |  |  |  |  |
| RRB | 4.32 | 2.67 |  |  |  |  |
| ADOS <sup>f</sup> |  |  |  |  |  |  |
| Social Affect | 6.04 | 2.59 |  |  |  |  |
| RRB | 4.73 | 2.78 |  |  |  |  |
| SRS T-score <sup>g</sup> | 70.59 | 12.06 | 47.24 | 8.79 |  |  |
| RBS <sup>h</sup> | 15.76 | 13.42 | 2.20 | 8.28 |  |  |
| SSP <sup>i</sup> | 138.62 | 27.28 | 175.97 | 16.18 |  |  |

TD, typically developing; SD, standard deviation; FSIQ, full-scale intelligence quotient; ADI, Autism Diagnostic Interview-Revised; RRB, restricted, repetitive behaviors; ADOS, Autism Diagnostic Observational Schedule 2; SRS, Social Responsiveness Scale 2nd Edition; RBS, Repetitive Behavior Scale-Revised; SSP, Short Sensory Profile.
<sup>a</sup> Statistical differences were assessed by two-sample *t*-test.
<sup>b</sup> In Schedule A, B, and C, there are 302 participants with autism and 229 participants with TD. Schedule D comprised 45 participants with autism and 23 TD individuals.
<sup>c</sup> Sex difference was examined by the chi-square test.
<sup>d</sup> ADHD rating scores were available for 500 participants, including 299 individuals with autism and 201 TD individuals.
<sup>e</sup> ADI scores were available for 332 participants.
<sup>f</sup> ADOS scores were available for 339 participants. We considered calibrated severity scores.
<sup>g</sup> Parent report SRS scores were available for 284 participants with autism and 135 TD individuals.
<sup>h</sup> RBS scores were available for 277 participants with autism and 133 TD individuals.
<sup>i</sup> SSP scores were available for 201 participants with autism and 115 TD individuals.
<sup>g, h, i</sup> In all questionnaires, the scores of the autism group only were used in our study, and they are all parent-rated.

### Clinical Measures

We used the Autism Diagnostic Interview-Revised (ADI) (25) and the Autism Diagnostic Observational Schedule 2 (ADOS) (26) to quantify past (ever and previous 4-to-5 years) and current autism symptoms of the social interaction, communication, and restricted repetitive behaviors (RRB) domains. We used T-scores (age- and sex-adjusted) of the Social Responsiveness Scale 2nd Edition (SRS) (27) in the autism group to assess severity of autistic traits/symptoms and the Repetitive Behavior Scale-Revised (RBS) (28) to measure repetitive and rigid behaviors associated with autism. Moreover, sensory processing abnormalities of autism were assessed with the Short Sensory Profile (SSP) (29). To examine associations between clinical features in autism and brain measures, we created two sets of clinical measures: 1) the subscale scores of ADI-R and ADOS, both instruments were rated by qualified examiners, and 2) the total scores of SRS, RBS, and SSP, we included parent-rated reports only and limited the analyses to within the autism group. Further, concerning the potential effect of comorbidity with Attention Deficit Hyperactivity Disorder (ADHD), we included comorbidity with ADHD as an additional covariate in analyses. ADHD symptoms were assessed with the ADHD DSM-5 rating scale that includes symptom scales of inattention and hyperactivity/impulsivity scores. The ADHD DSM-5 rating scale was based on either parent-report or self-report scores; self-report scores were only used when parent-reports were unavailable. The categorical output of the ADHD rating scale was used in this study. The summary for each of these clinical measures can be found in Table 1.

### MRI data acquisition

All participants were scanned on 3T MRI scanners (University of Cambridge: Siemens Verio; King’s College London: GE Medical Systems Discovery MR 750; Mannheim University: Siemens TimTrio; Radboud University: Siemens Skyra; Rome University: GE Medical Systems Sigma HDxTt; Utrecht University: Philips Medical Systems Achieva/Ingenia CX). High-resolution structural T1-weighted images were acquired with full head coverage, at 1.2 mm thickness with 1.2×1.2 mm in-plane resolution. For all other scanning parameters, please see Supplementary Table S1. Consistent image quality was ensured by a semi-automated quality control procedure.

### GM density estimation

Voxel-based morphometry (VBM) is a spatially-unbiased whole-brain approach that extracts voxel-wise GM density (the amount of GM at a voxel) estimations. We performed VBM analyses using the CAT12 toolbox (30) in SPM12 (Wellcome Department of Imaging Neuroscience, London, UK). T1-weighted images were automatically segmented into GM, white matter, and cerebrospinal fluid and affine registered to the MNI template to improve segmentation. All resulting segmented GM maps were then used to generate a study-specific template and registered to MNI space via a high-dimensional, nonlinear diffeomorphic registration algorithm (DARTEL) (31). A Jacobian modulation step was included using the flow fields to preserve voxel-wise information on local tissue volume. Images were smoothed with a 4 mm full-width half-max (FWHM) isotropic Gaussian kernel.

### Structural ICA decomposition

All participants’ VBM data were simultaneously decomposed into 100 spatially independent sources of spatial variation using MELODIC-ICA (15). Such ICA decomposition provides, at each independent component, a brain map reflecting a pattern of GM density covariation across participants, and a participant’s loading vector reflecting the contribution of each participant to each component. 100 components were chosen to capture as much variation as possible while remaining statistically powered (less than 25% of the total number of subjects (32)). However, the dependence on the model order (i.e., number of components) was also examined using different model orders; more precisely, in addition to the 100 dimensional factorization, we also considered an automatic dimension estimation approach as implemented in MELODIC-ICA and a 50 dimensional independent component factorization.

### Statistical approach

For completeness we first performed a standard mass-univariate statistical analyses directly on the GM densities. To that end we used a Generalized Linear Model (GLM) to detect group differences (autism vs. TD) on GM densities using the FMRIB Software Library v6.0 (FSL) (33). Participants’ VBM maps were considered as the dependent variable and diagnostic group as the independent factor, with age, sex, FSIQ, and scan site as covariates. Significance was assessed using permutation testing (5000 permutations) and correction for multiple comparisons was achieved using Threshold-Free Cluster Enhancement (TFCE, two-tailed, threshold at p<0.05) (34, 35).

Next, we considered the results of the ICA factorization of the VBM data. A GLM was used to examine differences between autistic and TD individuals, by using each participant’s loading to each component as a dependent variable, diagnostic group as independent variable, and age, sex, FSIQ, and scan site as regressors. To avoid the results of case-control differences being biased by IQ, we repeated the same procedure by excluding participants in Schedule D (FSIQ<75, more details see Supplementary subsection 2). Considering that autism is highly comorbid with ADHD (36), we also controlled for comorbidity with ADHD by adding a dummy-coded variable (with/without ADHD) to the original GLM analyses (autism vs. TD) in the case-control ICA analysis. Multiple comparison correction was implemented using false discovery rate (FDR) (p<0.05) (37).

We also explored independently the relationships between each estimated brain component and subscales of ADI and ADOS, SRS, RBS, and SSP in the autism group using GLM analyses and again correction for multiple comparisons was implemented with the FDR method (p<0.05). Then, to simultaneously explore the relationship between all the brain structural phenotypes estimated through ICA and all symptom phenotypes in the autism group, we used CCA. In the considered scenario, CCA is able to learn, at each canonical variate, linear projections of the brain structural sources and the behavioral measures that maximize the correlation between them at the participant level. Here, we performed two separate CCA analyses to link the independent components participants’ contributions to subsets of behavioral measures; in the first CCA analyses (CCA_1_) we included the subscales of ADI and ADOS as clinical measures and in the second (CCA_2_) we used total scores of SRS, RBS, and SSP. For each CCA analysis the statistical significance of each canonical variate was determined by permutation testing (10,000 permutations, Bonferroni corrected p<0.05/number of canonical variates). To evaluate the contribution of each independent source and each clinical measure to the CCA projections we used the structural coefficient of each variable as noted in (38). The reliability of the CCA results presented as well as its dependence on the number of subjects were tested using a leave-one-out cross-validation approach (Supplementary subsection 3).

## Results

### Mass-univariate statistics

The standard mass-univariate GLM analysis of the VBM data comparing cases and controls did not show significant group differences for voxel-wise GM volumes. Although no fully corrected statistical significance was assessed, we observed that the areas showing nominal significance (p<0.05) involved left temporal cortex, and bilateral cerebellum. We provide these uncorrected statistical results in supplementary materials Figure S2.

### Group effect on ICA decomposition

The structural data ICA decomposition provided a set of 100 independent spatial sources, each of which is connected to a vector that depicts the degree of each participant’s contribution to the corresponding components. For clarity, we further refer to these vectors as the participant loadings. Post-hoc GLM analyses of these participant loadings showed nominal significant case-control differences at nine independent components (ICs) (p<0.05, i.e. IC10, IC13, IC14, IC15, IC23, IC28, IC31, IC48, and IC 99, see Supplementary Figure S3). Of these, two components, IC10 (β=−0.147, p=8.850×10^−5^, effect size [Cohen’s d] d=−0.358) and IC14 (β=−0.132, p=5.450×10^−4^, d=−0.321), survived multiple comparison correction (FDR corrected, p<8.072×10^−4^). These results were not driven by age, sex, or scan site.

In Figure 1, we present summary images reflecting the brain areas involved in the structural variances occurring at these two components. The top row of Figure 1 shows that IC10 primarily relates to structural variation in the bilateral insula, inferior frontal gyrus (IFG), orbitofrontal cortex (OFC), and caudate nuclei. Among these brain regions, the bilateral caudate exhibits alterations in the opposite direction to the others. Given the negative beta coefficient obtained from the GLM analysis between participant loadings at IC10 and the diagnosis group labels, individuals with autism demonstrate increased GM densities in the bilateral caudate and decreased densities in the bilateral insula, IFG, and OFC. The bottom row of Figure 1 shows that IC14 mainly involves variations in the bilateral amygdala, hippocampus, and parahippocampal gyrus (PHG). Similarly, according to the sign of the beta values obtained through the GLM, the autism group shows decreased densities in the areas involved in IC14.

**Figure 1.**
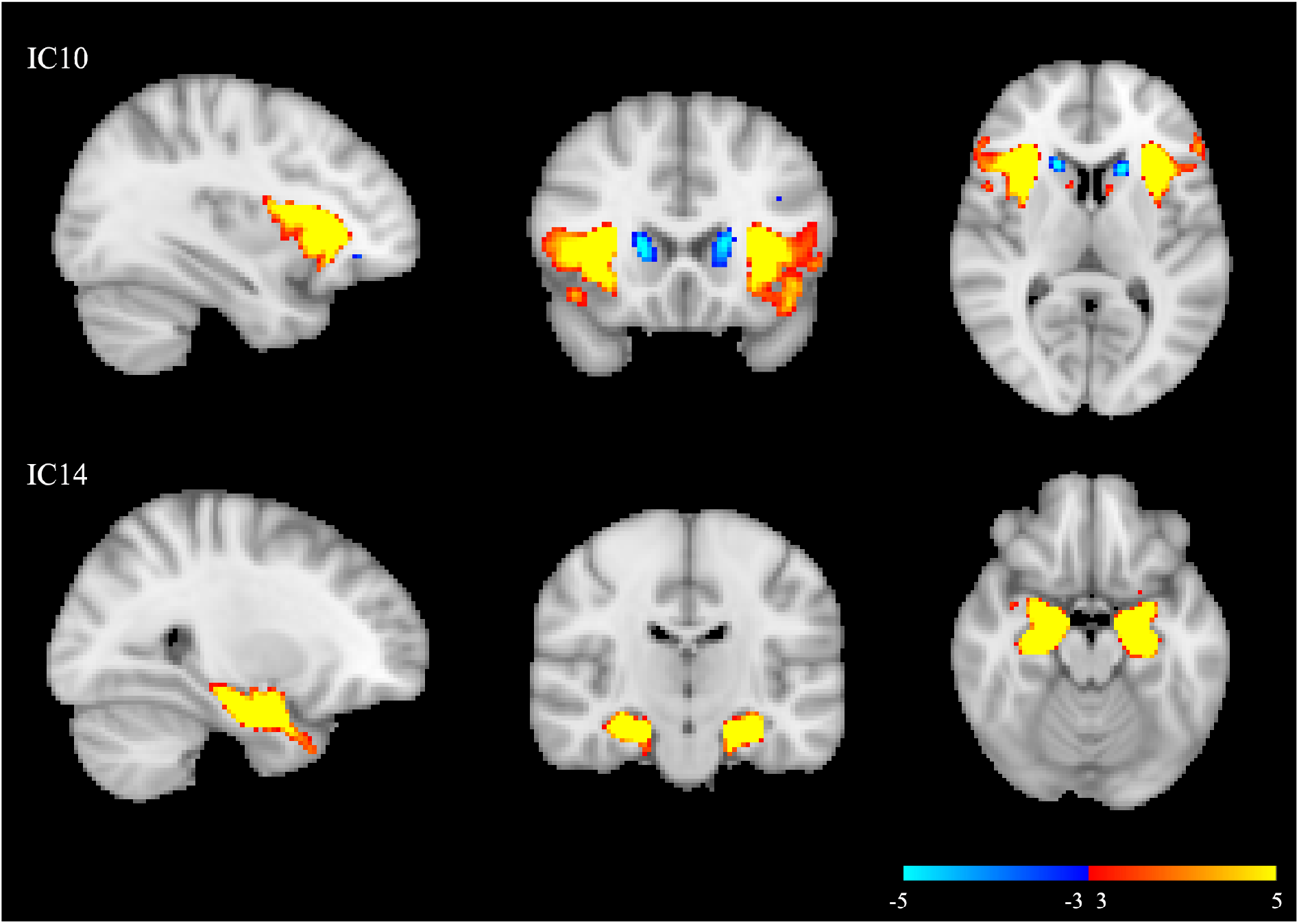
The components showed significant case-control differences. The component maps were thresholded at 3<|*Z*|<5. IC10, component number 10; IC14, component number 14.

The robustness of the ICA results to the model order choice was evaluated by considering, in addition to the original 100-dimensional factorization, an automatic dimensionality estimation procedure resulting in a 91-dimensional factorization, and a 50-dimensional factorization. We observed that the main components reported (IC10 and IC14) are highly reproducible independent of the model order choice. For details, see Supplementary materials subsection 6a.

To validate the ICA results not being biased by low IQ participants, an additional validation was performed by taking FSIQ into account to exclude the participants in Schedule D from the ICA factorization. This showed that a unique IC, corresponding to the original IC10, survived FDR correction (see Supplementary materials subsection 6b). Further, the effect of comorbidity with ADHD on brain structural variations was determined using data from 500 participants (for detailed demographic information, see Supplementary Table S4). This analysis showed that IC14 remained significantly associated with the autism group (FDR corrected, p=9.669 x10^−4^). However, IC10 was no longer associated with autism (p=0.004).

Further, post-hoc GLM analyses of the relationships between brain ICs and symptom ratings did not provide any significant associations (Supplementary Table S5).

### Relating gray matter spatial variation patterns to symptoms profiles

As a final step, we applied CCA to examine the associations between the 100 components and the two sets of clinical measures among the autism cases only. The CCA_1_ (linking ADI and ADOS subscale scores to brain measures), involved 325 autism participants and showed a Bonferroni corrected (p=0.05/5=0.010) significant relationship (Figure 2a,c, r=0.701, permutation p=0.008). In this main CCA mode, IC16, IC61, IC89, and IC14 were the highest contributors to the correlation with autism symptoms, and the ADOS RRB subscale loaded most on the association with the brain measures (Figure 3a, b). Among the four components, IC16 mainly involved density variations in bilateral thalamus and putamen (canonical weight: 0.447), IC61 in right lateral occipital and left superior parietal lobe (canonical weight: −0.366), and IC89 in the left precentral gyrus (canonical weight: −0.333). For details, see Supplementary Figure S6a. Note that IC14 is among the components previously reported showing linear significant case-control group effects. The regions involved in IC14 were mentioned above (canonical weight: −0.312). Since higher scores of the ADI and ADOS reflect more severe autism symptoms, positive values of IC16 suggest that higher loading on this component is related to more severe symptoms in autism, and negative values of IC61, IC89, and IC14 meant that lower loadings on these three ICs are associated with more severe symptoms. In Figure 2a, participants were color coded according to their ADOS-RRB scores to illustrate how the ADOS-RRB score drives the canonical correlation.

**Figure 2.**
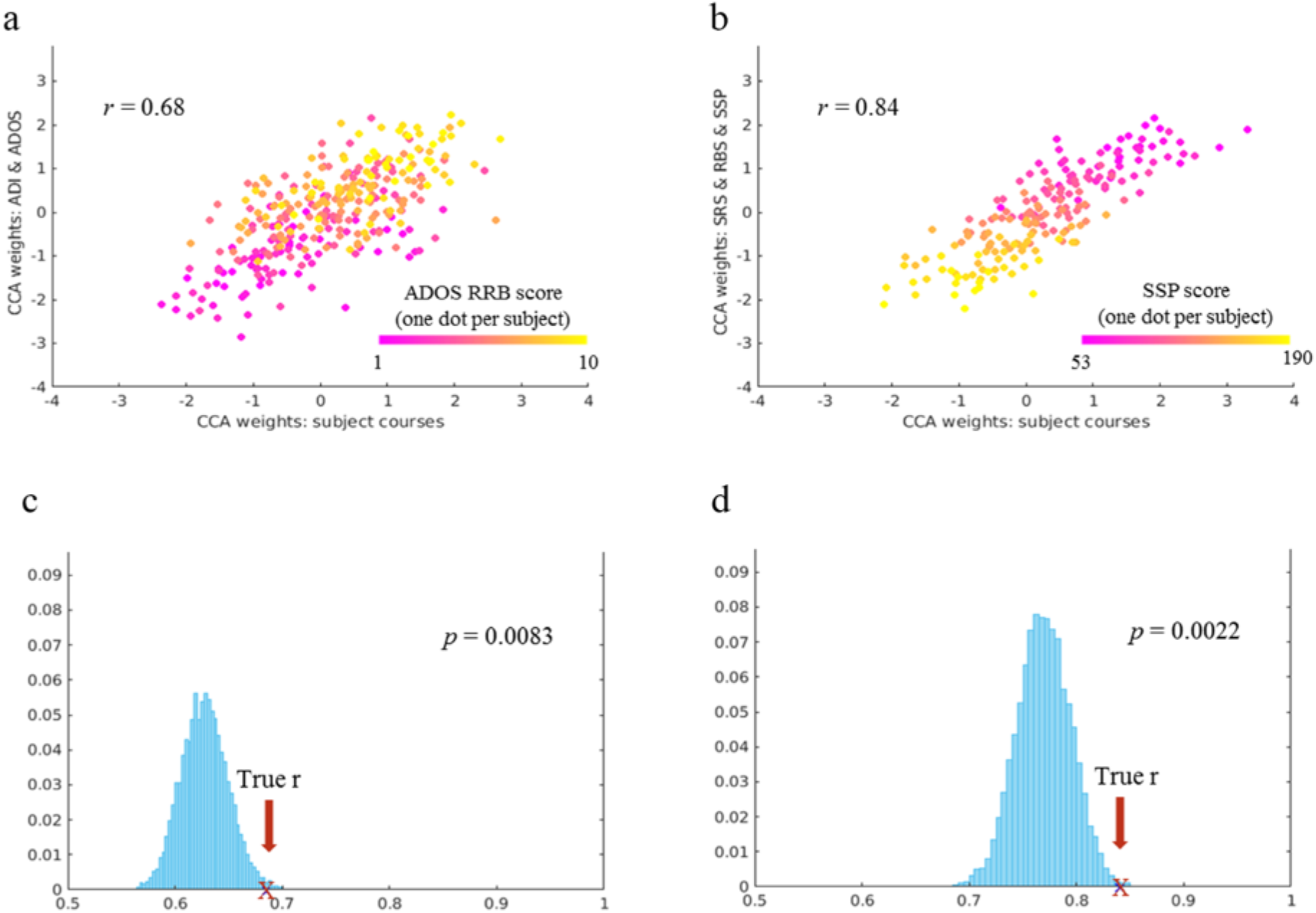
Main CCA mode scatter plots of participant loadings’ weights versus symptom profile weights. One dot per participant in each graph is coded with gradient colors according to the scores of ADOS RRB (a) and SSP (b), respectively. The second row shows the histograms of the null distribution of correlation values obtained from the main CCA mode at 10,000 random participants’ permutations in the autism sample with ADI and ADOS scores (c), and with SRS, RBS, and SSP scores (d). The true *r*-value is marked by a red cross. ADI, Autism Diagnostic Interview-Revised; ADOS, Autism Diagnostic Observational Schedule 2; SRS, Social Responsiveness Scale 2nd Edition; RBS, Repetitive Behavior Scale-Revised; SSP, Short Sensory Profile; RRB, restricted and repetitive behaviors.

**Figure 3.**
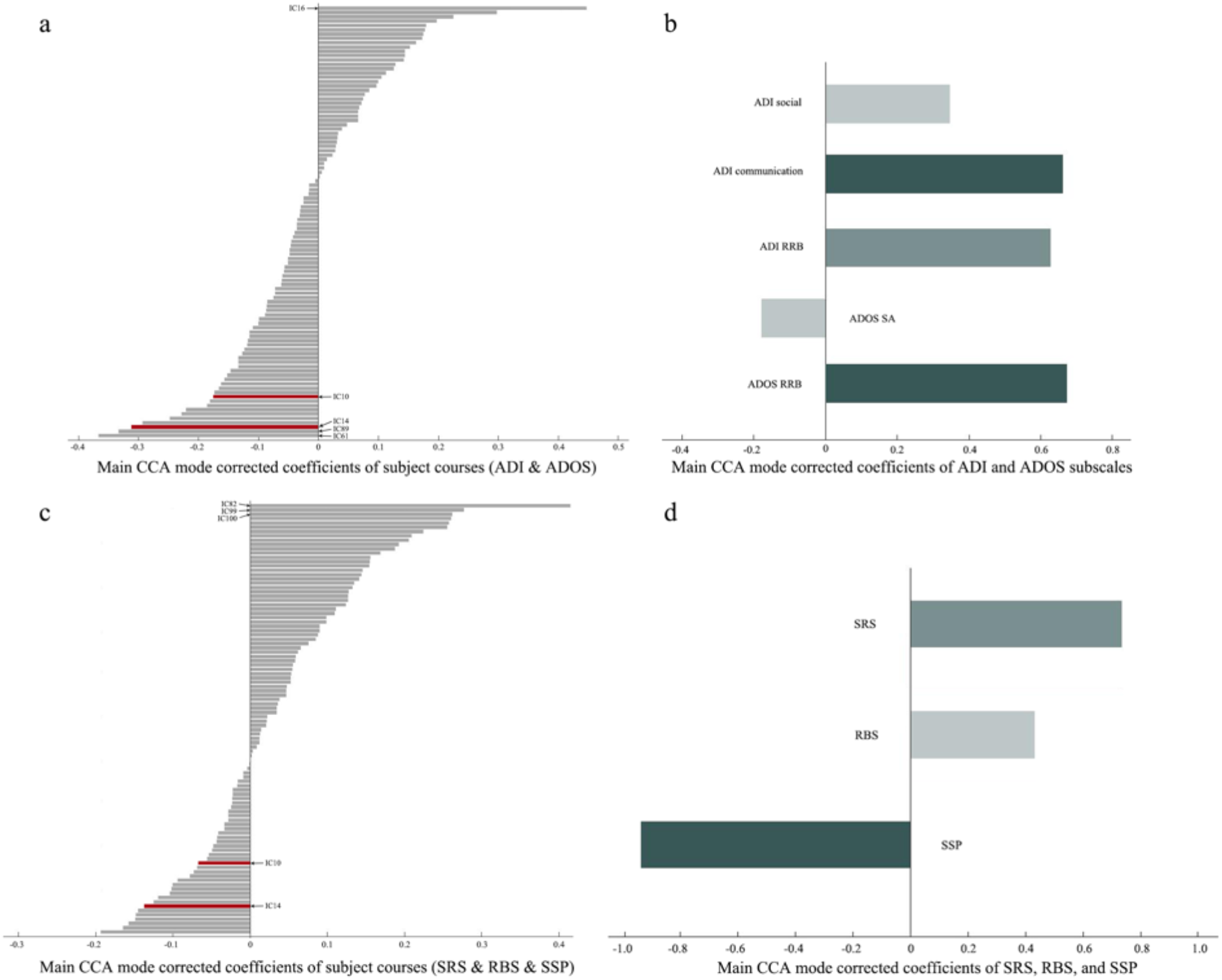
Main CCA mode corrected loadings of each component and symptom profiles. (a, c) display the degree that each component contributed to the main CCA mode related with ADI and ADOS (a), and with SRS, RBS, and SSP (c). The two components with significant group effects are displayed in red. (b, d) show the loadings of symptom profiles in each main CCA mode. CCA, canonical correlation analysis; ADI, Autism Diagnostic Interview-Revised; ADOS, Autism Diagnostic Observational Schedule 2; SRS, Social Responsiveness Scale 2nd Edition; RBS, Repetitive Behavior Scale-Revised; SSP, Short Sensory Profile.

In CCA_2_ we linked SRS, RBS, and SSP scores to the brain measures of 194 individuals with autism, which is 55.9% of all participants with autism (lower number due to missing questionnaire data). We found a Bonferroni corrected (p=0.05/3=0.017) significant relationship (Figure 2b, r=0.840, permutation p=0.002, Figure 2d). In this main CCA mode, IC82, IC99, and IC100 were the highest contributors to the correlation with behavior profiles, and SSP score loaded most on the association with the brain measures in the autism group (Figure 3c, d). IC82 mainly involved variations in the bilateral cerebellum (canonical weight: 0.414), IC99 in the left lateral occipital and parietal lobe, and bilateral precentral gyrus (canonical weight: 0.277), and IC100 in the left inferior frontal gyrus and right middle frontal lobe (canonical weight: 0.262). For details, see Supplementary Figure S6b. Similarly, lower loadings on these three ICs were related to more severe symptoms. In Figure 2b, each participant was color coded according to their SSP score, and it shows how SSP score drives the correlation. In this case, both IC10 and IC14 ranked outside the top 20 of the 100 components, suggesting that these two components with significant case-control difference have no strong contribution to the CCA_2_ correlation. However, for completeness, direct interpretation (referring to uncorrected coefficients) of the CCA_2_ weights ranks IC14 as the third strongest contributor to this canonical correlation (Supplementary Figure S7).

The CCA robustness analyses indicated that the main CCA modes of both CCA analyses were reliably estimated in a leave-one-subject out setting (Supplementary Figure S1). In CCA_1_, the weights of the main CCA mode of each leave-one-out analysis correlated on average above 0.94 with the weights of original main CCA mode in brain loadings and above 0.95 in behavior phenotypes when the sample was bigger than 122 subjects. In CCA_2_, the weights of the main CCA mode related on average above 0.92 in brain loadings and above 0.96 in behavior profiles when the sample was bigger than 111 subjects. Both CCA analyses are no reproducible for sample sizes smaller than (approximately) 100 subjects.

## Discussion

The present study used a reliable approach to quantify inter-individual differences in GM morphometry covariations in a deeply phenotyped large sample of individuals with and without autism. The standard, univariate VBM analysis did not show significant case-control differences. We then utilized an ICA decomposition of all participants GM density images, and similarly performed a case-control post-hoc statistical analyses. This analysis showed that autism was significantly associated with alterations in two independent sources of GM density covariations. These findings corroborated our hypothesis that alterations in brain morphometry in autism are associated with the combined influence of several regions rather than with a single region. In a following step, we applied CCA to explore multivariate associations between sets of continuous measures of core symptoms and sets of ICA-derived morphometry measures within the autism group, and were able to identify significant relationships between brain components and symptom profiles. Notably, one of the components which showed significant case-control differences was also among the highest loading components in the CCA.

Our findings showed two covarying sets of brain areas that structurally differed between cases and controls. While one source of GM density covariation, IC10, mainly related to the bilateral insula, IFG, OFG, and caudate, another source, IC14, primarily involved the bilateral amygdala, hippocampus, and PHG. The brain regions within each component are anatomically clustered and symmetrical, which indicates that the independent structural covariation alteration in the GM of individuals with autism is concentrated in nearby brain areas. This is in line with a previous study that used a similar approach (39). It is further in line with organizing principles of the brain that regions tend to be more interconnected when they are located close to each other (40, 41). However, when we compared the regions loading on the two components, the covarying regions of each component distribute in different brain locations. This suggests that neuroanatomic alterations underlying autism are more widely distributed at the whole brain level. It is of note that, when accounting for ADHD comorbidity, IC14 remained significant but IC10 did not. This suggests that IC14 is more specifically related to autism associated structural variations, even after linearly accounting for ADHD effects, while IC10 might reflect variations associated with both autism and ADHD.

The brain regions with high loadings on either of these two components, i.e. insula, amygdala, hippocampus and PHG have lower densities in autism and have earlier been associated with autism (9, 42). The opposite direction of the alteration of the caudate nucleus in autism has also previously been found (43). This is not the case for the IFG and OFG, which showed lower densities in autism in our study, where prior studies found mixed results (44, 45). Importantly, the brain regions identified by our analyses have earlier been implicated in the neurobiology and/or neurocognition of autism. In IC10, structural and/or functional alterations of the insula, IFG, and OFC have been associated with social and non-social cognitive impairments in autism (45–48). A meta-analysis reported abnormal functional activations of the insula, IFG, and OFG during social cognition tasks in autism (49). Additionally, variance of the caudate nucleus volume was found to correlate with the severity of RRB symptoms in autism (43). Together with deviant structural and functional connectivity between frontal cortical areas and striatum in autism (46, 50, 51), structural covariation in striatum and frontal areas may underlie atypical functional fronto-striatal connectivity, and this has been associated with repetitive behavior and executive functioning impairments in autism (3, 44). In the present study, the density of caudate nuclei increase simultaneously with densities decreasing on other areas in autism, which fits with the results of a few functional studies that indicate inverse functional changes of these areas (52). Particularly, the special pattern of GM densities changes in frontal and striatal areas might serve an important role in autism-related symptoms.

In IC14, we found decreased densities of amygdala and hippocampus, where the structural alterations have previously been related to social deficits in autism (6, 53). The amygdala, hippocampus, and PHG subserve cognitive and emotional functions that were found abnormal in individuals with autism (49, 54, 55). In addition to being involved in emotion and face processing, the three areas have been proposed as structures critical for working memory in autism (56). Furthermore, these cognitive domains exert bidirectional effects on each other, with atypical social-emotional processing influencing memory performance in individuals with autism, and memory being involved in complex information processing and executive functioning, which in turn affects social cognition (56, 57). In sum, given the potential functional interactions between these three brain areas, the structural covariance alterations of the amygdala, hippocampus, and PHG found in our study, may underlie or contribute to abnormal functional connections of these areas, and thus underlie poor performance on social cognition and memory tasks in individuals with autism.

Our multivariate correlation analyses moved from the case-control comparison to the use of continuous symptoms among individuals with autism and identified two prominent relationships between all structural brain covariances and symptoms in autism. Three of the four brain components that ranked top in this analysis did not show case-control differences, while there was one component (IC14) that differed between cases and controls and also significantly correlated with the severity of autism symptoms assessed by ADI and ADOS. The brain areas loading high on the brain components identified in the CCA are somewhat different from those implicated in the case-control analyses. These former brain areas are the thalamus, putamen, precentral gyrus, and parietal and occipital lobes in CCA_1_, and the cerebellum, frontal lobe, and again precentral gyrus, and parietal and occipital lobes in CCA_2_. These are foremost areas of the brain implicated in the processing and higher order integration of sensory information and motor functions. This makes sense since repetitive, rigid and stereotyped behaviors and abnormal sensory behaviors seem to drive the brain-behavior associations much more than the measures on social-communication symptoms. Note that variance within the different autism symptom domains (social-communication, repetitive behaviors and sensory abnormalities) was similar and cannot explain the differential symptom-brain associations.

Overall, the results of our multivariate analyses on case-control differences and on continuous measures of symptom severity among those with autism demonstrate the complexity of the brain morphometry correlates of autism. Brain areas involved are scattered across the whole brain and include the limbic system, basal ganglia, thalamus, cerebellum, precentral (motor) gyrus, and parts of the frontal, parietal, and occipital lobes.

## Strengths and limitations

The strengths of our study are the use of a large deeply phenotyped sample, bottom-up data-driven analyses, a multivariate approach for examining brain-symptom associations, and a large set of continuous measures of core autism symptoms. A limitation is that only 55.9% of the autism group had complete questionnaire data on continuous parent-reported symptom measures, which may have lowered statistical power for this analysis.

## Conclusions

We demonstrate brain morphometry differences between individuals with autism and typical controls in the inter-regional covariation of the insula, frontal area, caudate, amygdala, hippocampus, and PHG. Further, we highlight associations between covariation in density of the thalamus, putamen, precentral gyrus, frontal, parietal, and occipital lobes, and the cerebellum, and core autism symptoms, in particular repetitive behaviors and abnormal sensory behavior. Future studies may link our morphometry findings with data on brain function obtained from cognitive tests and/or functional and resting-state MRI, and with genomics data.

## Abbreviations

ADI: Autism Diagnostic Interview-Revised
ADOS: Autism Diagnostic Observational Schedule 2
CCA: Canonical Correlation Analysis
DSM-IV: Diagnostic and Statistical Manual-IV
EU-AIMS: European Autism Interventions—A Multicentre Study for Developing New Medications
FDR: false discovery rate
FSIQ: full-scale intelligence quotient
GLM: Generalized Linear Model
GM: gray matter
IC: Independent Component
ICA: Independent Component Analysis
ICD-10: International Statistical Classification of Diseases and Related Health Problems 10th Revision
ID: intellectual disability
IFG: inferior frontal gyrus
OFC: orbitofrontal cortex
MRI: Magnetic resonance imaging
PHG: parahippocampal gyrus
RBS: Repetitive Behavior Scale-Revised
RRB: Restricted and Repetitive Behaviors
SA: social affect
SD: standard deviation
SRS: Social Responsiveness Scale 2nd Edition
SSP: Short Sensory Profile
TD: typically developing
TFCE: Threshold-Free Cluster Enhancement
VBM: Voxel-based Morphometry

## Ethics approval and consent to participate

Ethical approval for this study was obtained through ethics committees at each site.

## Consent for publication

Consent for publication was obtained from all participants prior to the study.

## Availability of data and materials

Data collected in EU-AIMS LEAP are stored and curated at the central EU-AIMS database at the Pasteur Institute in Paris. The database is open to members of the wider scientific community upon request and submission of a paper and data analytic proposal.

## Competing interests

JKB has been a consultant to, advisory board member of, and a speaker for Janssen Cilag BV, Eli Lilly, Shire, Lundbeck, Roche, and Servier. He is not an employee of any of these companies, and not a stock shareholder of any of these companies. He has no other financial or material support, including expert testimony, patents or royalties. CFB is director and shareholder in SBGNeuro Ltd. TB served in an advisory or consultancy role for Lundbeck, Medice, Neurim Pharmaceuticals, Oberberg GmbH, Shire, and Infectopharm. He received conference support or speaker’s fee by Lilly, Medice, and Shire. He received royalities from Hogrefe, Kohlhammer, CIP Medien, and Oxford University Press. TC has received consultancy from Roche and received book royalties from Guildford Press and Sage. DGM has been a consultant to, and advisory board member, for Roche and Servier. He is not an employee of any of these companies, and not a stock shareholder of any of these companies. The present work is unrelated to the above grants and relationships. The other authors report no biomedical financial interests or potential conflicts of interest.

## Funding

The work is supported by the European Union Seventh Framework Programme Grant Nos. 602805 (AGGRESSOTYPE) (to JKB), 603016 (MATRICS) (to JKB), and 278948 (TACTICS) (to JKB); European Community’s Horizon 2020 Programme (H2020/2014-2020) Grant Nos. 643051 (MiND) (to JKB) and 642996 (BRAINVIEW) (to JKB); the Netherlands Organization for Scientific Research VIDI Grant Nos. 864.12.003 (to CFB); Wellcome Trust UK Strategic Award Grant No.098369/Z/12/Z (to CFB); the Autism Research Trust (to SBC); and EU-AIMS (European Autism Interventions), which receives support from Innovative Medicines Initiative Joint Undertaking Grant No.115300, the resources of which are composed of financial contributions from the European Union’s Seventh Framework Programme (Grant No. FP7/2007-2013), from the European Federation of Pharmaceutical Industries and Associations companies’ in-kind contributions; and AIMS-2-TRIALS (Autism Innovative Medicine Studies-2-Trials), which has received funding from the Innovative Medicines Initiative 2 Joint Undertaking under grant agreement No. 777394, and this Joint Undertaking receives support from the European Union’s Horizon 2020 research and innovation programme and EFPIA and AUTISM SPEAKS, Autistica, SFARI.

## Authors’ contributions

JT, SD, CM, TB, RH, SB-C, AR, EL, TC, DGM, CE, CFB, JKB designed the study, developed data acquisition and/or analysis protocols. TM ran analyses, visualized the findings, wrote the first draft, and revised the draft; AL conceptualized and supervised the analysis, and revised the manuscript; DLF ran the VBM analysis and revised the draft; JT & NJF revised the draft; CFB supervised and revised the manuscript; JKB supervised and revised the manuscript. All authors read and approved the final manuscript.

## Acknowledgements

We thank all participants and their families for participating in this study. We gratefully acknowledge the contributions of all members of the EU-AIMS LEAP group: Jumana Ahmad, Sara Ambrosino, Bonnie Auyeung, Tobias Banaschewski, Simon Baron-Cohen, Sarah Baumeister, Christian F. Beckmann, Sven Bölte, Thomas Bourgeron, Carsten Bours, Michael Brammer, Daniel Brandeis, Claudia Brogna, Yvette de Bruijn, Jan K. Buitelaar, Bhismadev Chakrabarti, Tony Charman, Ineke Cornelissen, Daisy Crawley, Flavio Dell’Acqua, Guillaume Dumas, Sarah Durston, Christine Ecker, Jessica Faulkner, Vincent Frouin, Pilar Garcés, David Goyard, Lindsay Ham, Hannah Hayward, Joerg Hipp, Rosemary Holt, Mark H. Johnson, Emily J.H. Jones, Prantik Kundu, Meng-Chuan Lai, Xavier Liogier D’ardhuy, Michael V. Lombardo, Eva Loth, David J. Lythgoe, René Mandl, Andre Marquand, Luke Mason, Maarten Mennes, Andreas Meyer-Lindenberg, Carolin Moessnang, Nico Mueller, Declan G.M. Murphy, Bethany Oakley, Laurence O’Dwyer, Marianne Oldehinkel, Bob Oranje, Gahan Pandina, Antonio M. Persico, Annika Rausch, Barbara Ruggeri, Amber Ruigrok, Jessica Sabet, Roberto Sacco, Antonia San José Cáceres, Emily Simonoff, Will Spooren, Julian Tillmann, Roberto Toro, Heike Tost, Jack Waldman, Steve C.R. Williams, Caroline Wooldridge, and Marcel P. Zwiers. TM is funded by the PhD scholarship (201806010408) of the Chinese Scholarship Council (CSC).

