## Supplementary Materials for "Gray matter covariations and core symptoms of autism. The EU-AIMS Longitudinal European Autism Project"

**Supplementary Information**

### 1. Acquisition parameters

**Table S1**. Summary of acquisition parameters across sites

| Site | Manufacturer | Model | Software Version | Acquisition sequence | Coverage | Slices | Thickness [mm] | Resolution [mm$^3$] | TR [s] | TE [ms] | FA [°] | FOV |
|---|---|---|---|---|---|---|---|---|---|---|---|---|
| Cambridge | Siemens | Verio | Syngo MR B17 | Tfl3d1_ns | 256*256 | 176 | 1.2 | 1.1*1.1*1.2 | 2.3 | 2.95 | 9 | 270 |
| London | GE Medical systems | Discovery mr750 | LX MR DV23.1_V02 _1317.c | SAG ADNI GO ACC SPGR | 256*256 | 196 | 1.2 | 1.1*1.1*1.2 | 7.31 | 3.02 | 11 | 270 |
| Mannheim | Siemens | TimTrio | Syngo MR B17 | MPRAGE ADNI | 256*256 | 176 | 1.2 | 1.1*1.1*1.2 | 2.3 | 2.93 | 9 | 270 |
| Nijmegen | Siemens | Skyra | Syngo MRD13 | Tfl3d1_16ns | 256*256 | 176 | 1.2 | 1.1*1.1*1.2 | 2.3 | 2.93 | 9 | 270 |
| Rome | GE Medical systems | Signa HDxt | 24/LX/MR HD16.0_V02 _1131.a | SAG ADNI GO ACC SPGR | 256*256 | 172 | 1.2 | 1.1*1.1*1.2 | 5.96 | 1.76 | 11 | 270 |
| Utrecht | Philips Medical Systems | Achieva /Ingenia CX | 3.2.3, 3.2.3.1 | ADNI GO 2 | 256*256 | 170 | 1.2 | 1.1*1.1*1.2 | 6.76 | 3.1 | 9 | 270 |

TR: repetition time; TE: echo time; FA: flip angle; FOV: field of view.

### 2. Demographic information of each schedule

Participants were split into four schedules depending on their age and full-scale intelligence quotient (FSIQ). Schedule A included adults aged 18-30 years, Schedule B included adolescents aged 12-17 years, and Schedule C included children aged 6-11 years. In schedules A-C, all participants had a FSIQ in the typical range (FSIQ≥75). Lastly, Schedule D comprised adolescents and adults aged 12-30 years with mild intellectual disability (ID) (50≤FSIQ<75) (Table S2).

**Table S2**. Demographic information of participants in each schedule

| | Schedule A | | Schedule B | | Schedule C | | Schedule D | |
|---|---|---|---|---|---|---|---|---|
| Variable | Autism | TD | Autism | TD | Autism | TD | Autism | TD |
| N | 114 | 85 | 115 | 85 | 73 | 59 | 45 | 23 |
| Age,mean,[SD] | 22.54 [3.25] | 22.99 [3.34] | 14.88 [1.77] | 15.38 [1.74] | 9.60 [1.44] | 9.74 [1.49] | 18.77 [4.41] | 18.58 [4.39] |
| IQ,mean,[SD] | | | | | | | | |
| Full-scale IQ | 104 [15] | 109 [13] | 103 [15] | 106 [13] | 107 [14] | 113 [13] | 67 [5] | 64 [9] |
| Performance IQ [a] | 106 [16] | 108 [15] | 105 [18] | 107 [15] | 106 [14] | 112 [15] | 65 [10] | 64 [11] |
| Verbal IQ [a] | 103 [16] | 109 [15] | 100 [15] | 104 [14] | 107 [15] | 113 [14] | 69 [9] | 64 [11] |
| Sex, N, [%] | | | | | | | | |
| Male | 82 [71.93] | 56 [65.88] | 90 [78.26] | 59 [69.41] | 52 [71.23] | 36 [61.02] | 29 [64.44] | 12 [52.17] |
| Female | 32 [28.07] | 29 [34.12] | 25 [21.74] | 26 [30.59] | 21 [28.77] | 23 [38.98] | 16 [35.56] | 11 [47.83] |
| Site, N | | | | | | | | |
| Cambridge | 17 | 10 | 19 | 9 | 12 | 10 | 3 | 0 |
| KCL | 52 | 39 | 36 | 18 | 24 | 8 | 21 | 13 |
| Mannheim | 5 | 5 | 19 | 24 | 2 | 7 | 2 | 0 |
| Nijmegen | 25 | 12 | 29 | 26 | 27 | 22 | 19 | 10 |
| Utrecht | 16 | 19 | 12 | 9 | 9 | 13 | 1 | 0 |

TD, typically developed; SD, standard deviation; IQ, intelligence quotient.

[a] In Schedule C, there were 3 individuals with autism missing the performance and verbal IQ data.

### 3. Reproducibility of CCA results

To assess the reproducibility of CCA results, we employed a leave-one-out (LOO) approach by randomly resampling subsets of the sample with participant number from 50 to 325 (ADI&ADOS)/194 (SRS&RBS&SSP), and repeating each LOO analysis 50 times. In each subset, we separately correlated the main mode weights of brain loadings and behavior profiles, which were generated from LOO analysis of CCA, with the weights of the original main mode. We then used the mean and standard deviation of r values to evaluate the reproducibility of CCA in different sample sizes. In $CCA_1$ (ADI&ADOS), the weights of the main CCA mode of each leave-one-out analysis correlated on average above 0.94 with the weights of original main CCA mode in brain loadings and above 0.95 in behavior profiles when the sample was bigger than 122. In $CCA_2$ (SRS&RBS&SSP), the weights of the main CCA mode related on average above 0.92 in brain loadings and above 0.96 in behavior profiles when the sample was bigger than 111. Both CCA analyses are no reproducible for sample sizes smaller than (approximately) 100 subjects. (Figure S1).

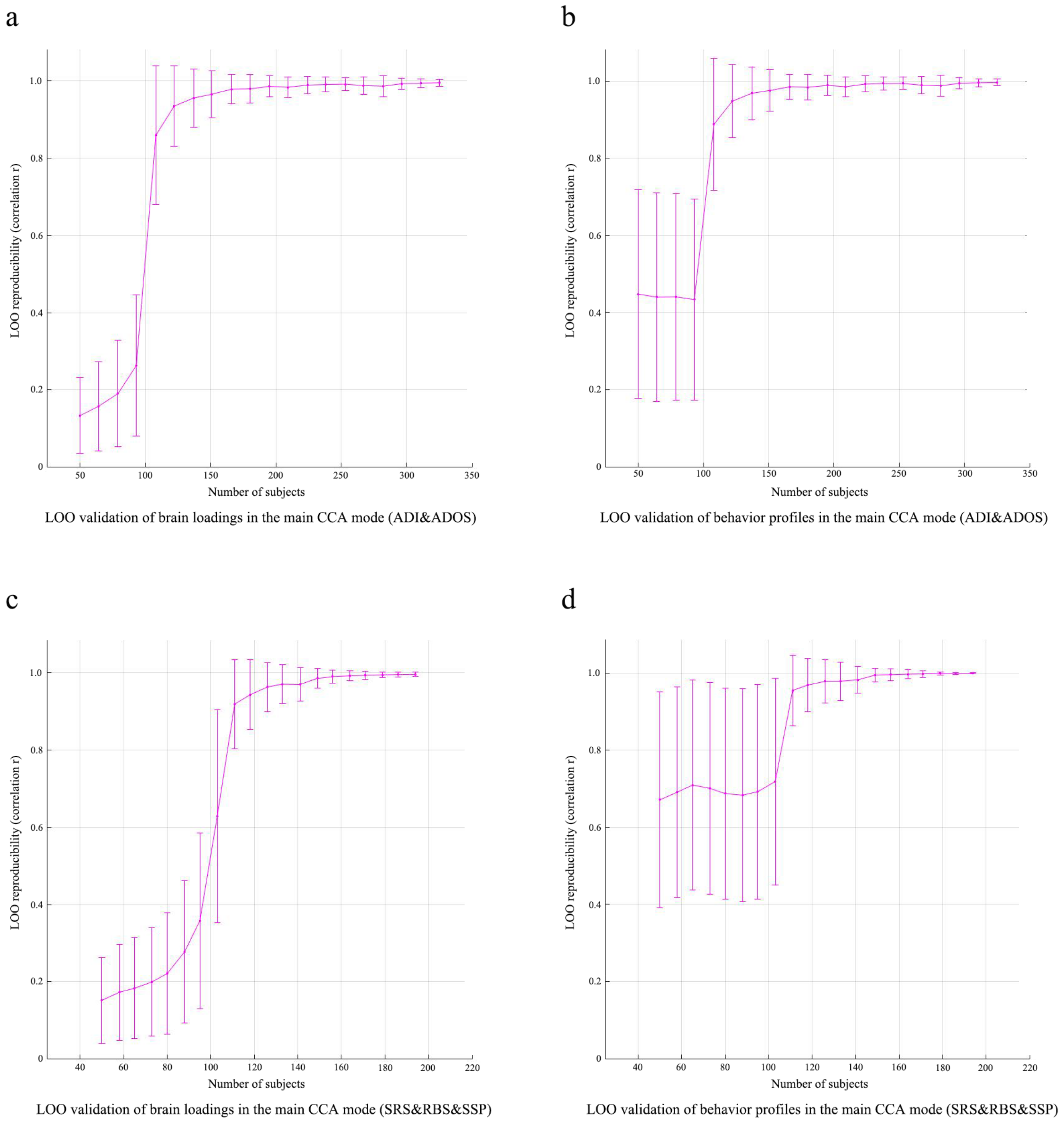

**Figure S1**. LOO validation of the main CCA modes in both CCA. (a, c) display the reproducibility of brain components of the main CCA mode related with ADI and ADOS (a), and with SRS, RBS, and SSP (c). (b, d) show the reproducibility of symptom profiles in each main CCA mode. LOO, leave-one-out; CCA, canonical correlation analysis; ADI, Autism Diagnostic Interview-Revised; ADOS, Autism Diagnostic Observational Schedule 2; SRS, Social Responsiveness Scale 2nd Edition; RBS, Repetitive Behavior Scale-Revised; SSP, Short Sensory Profile.

### 4. Group differences at voxel-wise gray matter volumes

The standard mass-univariate GLM analysis of the VBM data comparing cases and controls did not show significant group differences for voxel-wise GM densities. Figure S2 presents each voxel t-statistics and it is thresholded at uncorrected p<0.05. The regions showing nominal significance involve the left temporal cortex and bilateral cerebellum.

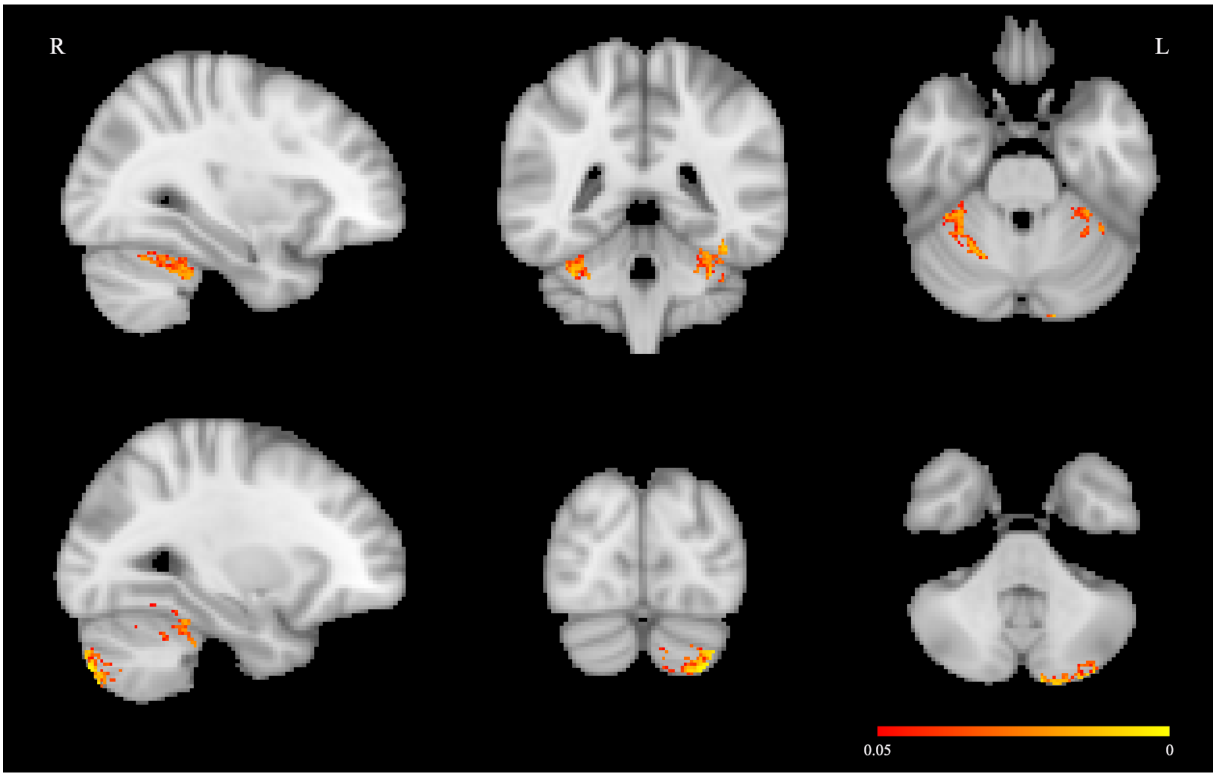

**Figure S2**. Results of case-control differences of voxel-wise gray matter densities (p<0.05, uncorrected).

### 5. Independent components with case-control differences

Nine independent components (ICs) showed nominal significant case-control differences ($p<0.05$, i.e. IC10, IC13, IC14, IC15, IC23, IC28, IC31, IC48, and IC 99, Figure S1), among which IC10 ($\beta$=-0.175, $p=8.850\times10^{-5}$) and IC14 ($\beta$=-0.152, $p=5.450\times10^{-4}$) survived FDR correction ($p<8.072\text{x}10^{-4}$).

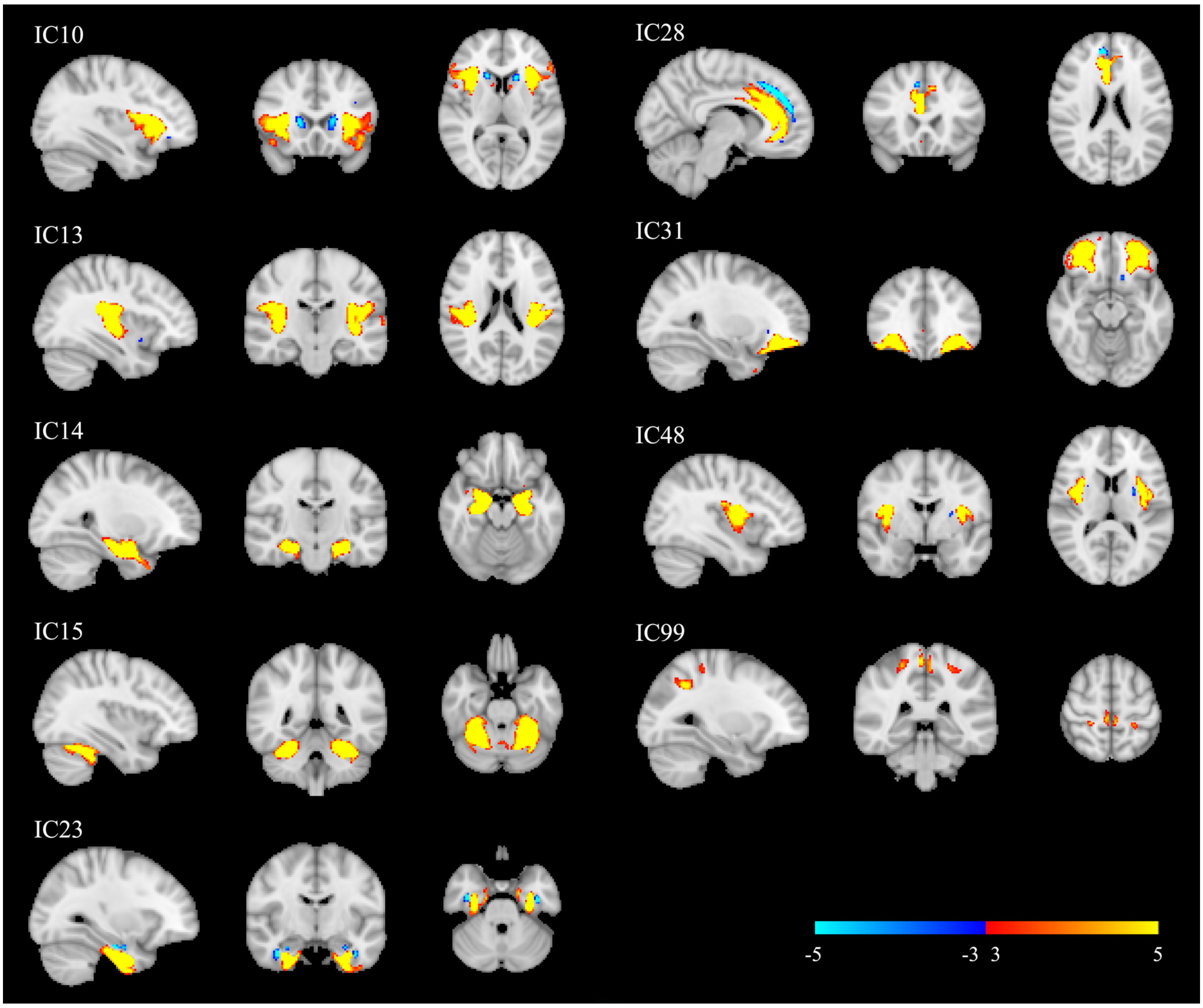

**Figure S3**. The components showed significant case-control differences ($p<0.05$, uncorrected). The component maps were thresholded at $3<|Z|<5$. IC, independent component.

### 6. Robustness assessment of the ICA model

To assess the reproducibility of components IC10 and IC14 obtained from the mainly reported 100-dimensional decomposition, we first correlated the participant loadings from the 100-dimensional factorization with the participant loadings obtained from an alternative factorization. This allowed us to identify the two components from the alternative factorization that are more strongly correlated to IC10 and IC14 respectively, and evaluate then the spatial reproducibility achieved at the alternative factorization.

#### a. Results validated by automatic dimension estimation and 50-dimensional factorization

Post-hoc statistics, for automatic estimation (91 ICs) and the 50-dimensional IC analysis factorization showed that the composition of components with significant group effects were similar to the original analysis with 100 components. Significant results of both dimensional factorizations are almost equivalent to the IC10 (automatic dimension: $p= 2.109\times10^{-4}$ , 50 dimension: $p=0.002$) and IC15 (automatic dimension: $p=3.557\times10^{-4}$, 50 dimension: $p=2.778\times10^{-4}$) in the analysis of 100-dimensional factorization, however, the ICs corresponding to IC14 in automatic dimension ($p=4.733\times10^{-4}$) and 50 dimensions ($p=0.003$) did not survive FDR correction (automatic dimension: $p<4.733\times10^{-4}$; 50 dimension factorization: $p<0.003$) (Figure S4).

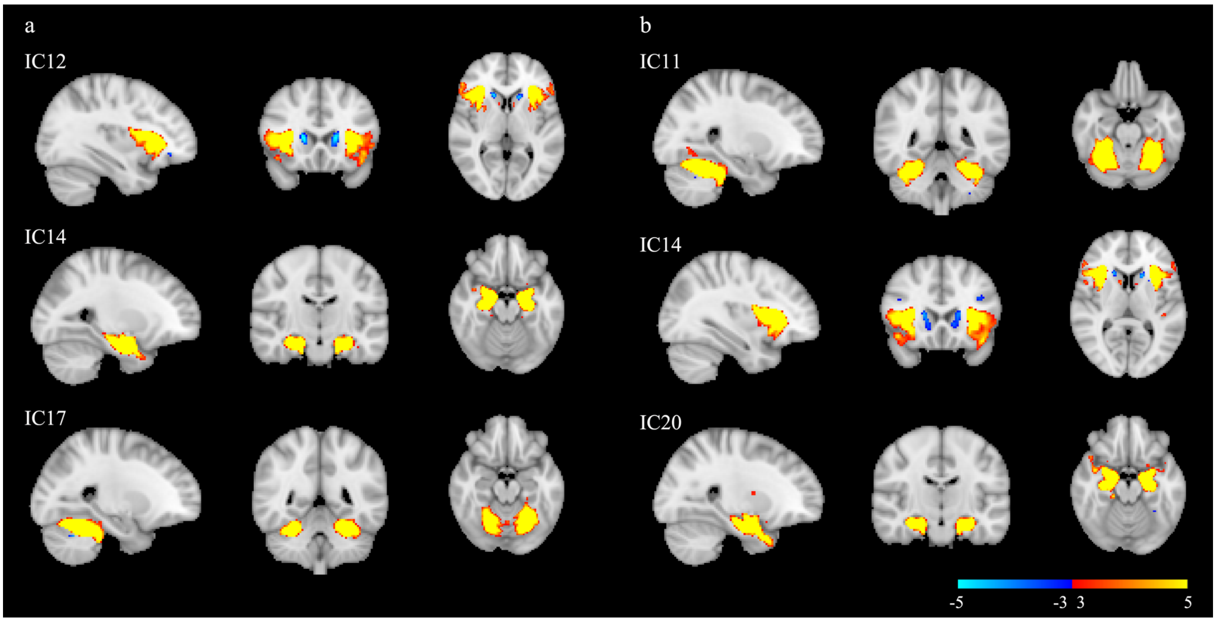

**Figure S4**. The components showed significant case-control differences. Panel **a** shows the components in **automatic** dimensional factorization. IC12 ($p=2.109\times10^{-4}$), corresponding to IC10 in 100-dimensional factorization, and IC17 ($p=3.557\times10^{-4}$) survived multiple comparison correction ($p<4.733\times10^{-4}$). IC14 ($p=4.733\times10^{-4}$), corresponding to the IC14 in 100-dimensional factorization did not survive correction. Panel **b** shows the components in 50-dimensional factorization. IC11 ($p=2.778\times10^{-4}$), and IC14 ($p=0.002$), corresponding to IC10 in 100-dimensional factorization, survived multiple comparison correction ($p<0.003$). IC20 ($p=0.003$), corresponding to IC14 in 100-dimensional factorization did not survive correction. The component maps were thresholded at $3<|Z|<5$. IC, independent component.

**Table S3**. Summary of robustness assessment of ICA results (correlation results)

| | conditions | corresponding IC | participant loadings | | spatial maps | |
|---|---|---|---|---|---|---|
| | | | r | p | r | p |
| IC10 | automatic dimension | IC12 | 0.990 | p<0.001 | 0.979 | p<0.001 |
| | 50 dimensions | IC14 | 0.941 | p<0.001 | 0.879 | p<0.001 |
| IC14 | automatic dimension | IC14 | 0.994 | p<0.001 | 0.990 | p<0.001 |
| | 50 dimensions | IC20 | 0.927 | p<0.001 | 0.870 | p<0.001 |

IC, independent component.

**b. Results validated by excluding intellectual disability participants**

To validate the results not being biased by low IQ participants we excluded the participants in Schedule D (FIQ < 75) and performed an analogous 100-dimensional ICA decomposition, and automatic dimensional estimated factorizations (85 ICs) followed by post-hoc statistics again. In the 100-dimensional factorization case, we found the components with significant group effect were equivalent to IC10 and IC14 in the original analysis with the Schedule D participants included (Figure S5a), however, they did not survive FDR correction ($p<1.600\times10^{-5}$). Similarly, in the automatic dimensional factorization case, we found significant ICs corresponding to IC10 and IC14, however the IC corresponding to IC14 did not survive FDR correction ($p<7.531\times10^{-4}$) (Figure S5b).

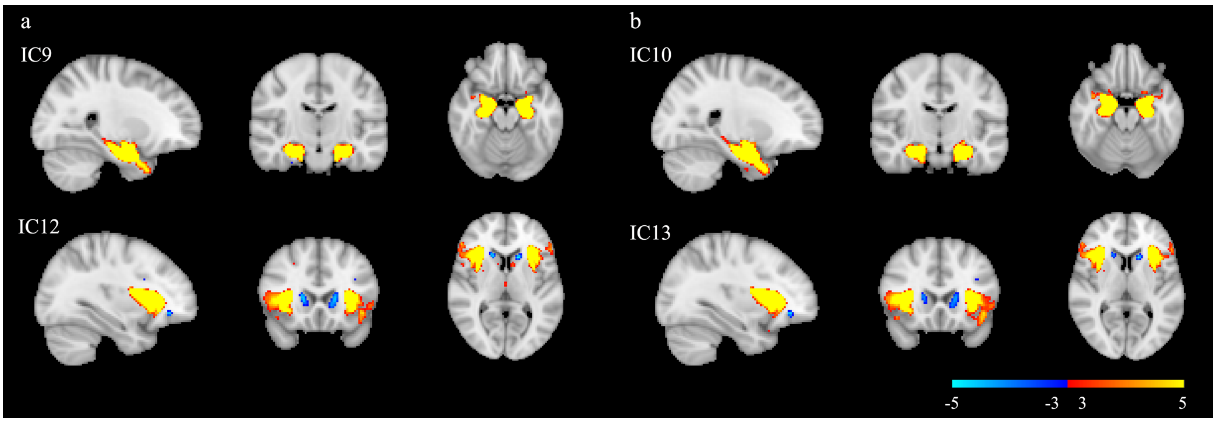

**Figure S5**. The components showed significant case-control differences. Panel **a** shows the components in **100** dimensional factorization **excluding** Schedule D participants. IC9 corresponds to IC14 in the 100 dimensional factorization including Schedule D participants (p=0.001), and IC12 corresponds to IC10 ($p=1.600\times10^{-5}$). Neither of these ICs survive FDR correction ($p<1.600\times10^{-5}$). Panel **b** shows the components in **automatic** dimensional factorization **excluding** Schedule D participants. IC10 corresponds to IC14 in 100 dimensional factorization including Schedule D participants ($p=7.531\times10^{-4}$), and IC13 ($p=3.830\times10^{-5}$) corresponds to IC10 that survived FDR correction ($p<7.531\times10^{-4}$). The component maps were thresholded at $3<|Z|<5$. IC, independent component.

### 7. Demographic information of the 500 participants used to analyze the effect of comorbidity

**Table S4**. Demographic information of the 500 participants used to analyze the effect of comorbidity

| **Demographic** | Autism, n = 299 | | TD, n = 201 | | $t/\chi^2$ | p value |
|---|---|---|---|---|---|---|
| | Mean | SD | Mean | SD | | |
| Age, years [a] | 16.79 | 5.59 | 16.77 | 5.59 | -0.037 | 0.970 |
| FIQ [a] | 99.74 | 19.05 | 105.31 | 17.26 | 3.397 | p=0.001 |
| | n | % | n | % | | |
| Sex, male/female [b] | 212/87 | 70.9/29.1 | 127/74 | 63.2/36.8 | 3.280 | 0.070 |
| **Clinical** | n | % | n | % | | |
| ADHD rating scale [b], with ADHD/without | 139/160 | 46.5/53.5 | 21/180 | 10.4/89.6 | 71.750 | p<0.001 |

TD, typically developed; SD, standard deviation; FIQ, full-scale intelligence quotient; ADHD, attention deficit hyperactivity disorder.

[a] Statistical differences were assessed by two sample *t*-test.

[b] Statistical difference was examined by the chi-square test.

### 8. GLM results of the association between brain components and symptom profiles

**Table S5**. GLM results of the association between brain components and symptom profiles in autism group (uncorrected, p<0.05)

| component | ADI | | | | | | ADOS | | | | SRS [a] | | RBS | | SSP | |
|---|---|---|---|---|---|---|---|---|---|---|---|---|---|---|---|---|
| | social | | communication | | RRB | | social affect | | RRB | | | | | | | |
| | b | p | b | p | b | p | b | p | b | p | b | p | b | p | b | p |
| 2 | -0.150 | 0.009 | | | | | | | | | | | | | | |
| 3 | | | | | | | -0.194 | $6.169 \times 10^{-4}$ | | | | | | | | |
| 5 | | | | | | | -0.147 | 0.007 | | | | | | | | |
| 6 | -0.120 | 0.046 | | | | | | | | | | | | | | |
| 9 | | | | | 0.120 | 0.032 | | | | | | | | | | |
| 12 | -0.124 | 0.016 | | | | | -0.112 | 0.033 | | | | | | | | |
| 14 | | | | | | | | | -0.133 | 0.022 | | | | | | |
| 15 | | | | | | | | | 0.107 | 0.049 | | | | | | |
| 21 | | | 0.108 | 0.048 | | | | | | | | | | | | |
| 24 | | | | | | | | | 0.114 | 0.035 | | | | | | |
| 27 | | | | | | | | | | | 0.116 | 0.030 | | | | |
| 31 | | | | | | | | | -0.130 | 0.010 | | | | | | |
| 33 | -0.108 | 0.039 | | | | | | | | | | | | | | |
| 40 | | | | | | | | | | | | | 0.110 | 0.035 | | |
| 41 | -0.131 | 0.016 | | | | | -0.171 | 0.002 | | | -0.121 | 0.031 | | | | |
| 42 | | | | | | | | | | | 0.108 | 0.046 | | | | |
| 44 | 0.141 | 0.006 | 0.110 | 0.029 | | | | | | | | | | | | |
| 51 | | | | | | | | | | | | | | | 0.130 | 0.044 |
| 57 | | | | | -0.123 | 0.048 | | | | | | | | | | |
| 59 | | | | | | | | | 0.167 | 0.001 | | | | | | |
| 61 | | | | | -0.120 | 0.023 | | | | | | | | | | |
| 63 | | | | | | | -0.138 | 0.028 | | | | | | | | |

**Table S5**. GLM results of the association between brain components and symptom profiles in autism group (*p*<0.05, continued)

| component | ADI | | | | | | ADOS | | | | SRS [a] | | RBS | | SSP | |
|---|---|---|---|---|---|---|---|---|---|---|---|---|---|---|---|---|
| | social | | communication | | RRB | | social affect | | RRB | | | | | | | |
| | b | p | b | p | b | p | b | p | b | p | b | p | b | p | b | p |
| 65 | | | | | -0.112 | 0.039 | | | | | | | | | | |
| 69 | | | | | | | | | | | | | | | -0.142 | 0.022 |
| 82 | | | | | | | | | | | | | | | -0.212 | 4.169x10-4 |
| 89 | | | -0.108 | 0.038 | | | | | | | | | | | | |
| 90 | | | | | | | | | | | | | -0.125 | 0.032 | | |
| 95 | -0.131 | 0.015 | | | | | | | | | | | | | | |
| 97 | | | | | | | | | | | | | | | -0.125 | 0.038 |
| 98 | | | | | | | | | | | | | | | -0.127 | 0.048 |
| 100 | | | | | | | | | | | | | 0.112 | 0.034 | -0.170 | 0.005 |

The association analyses were only performed in autism group. ADI, Autism Diagnostic Interview-Revised; RRB, restricted, repetitive behaviors; ADOS, Autism Diagnostic Observational Schedule 2; SRS, Social Responsiveness Scale 2nd Edition; RBS, Repetitive Behavior Scale-Revised; SSP, Short Sensory Profile.

[a] We used SRS parent T-scores.

### 9. Components with highest loadings in CCA

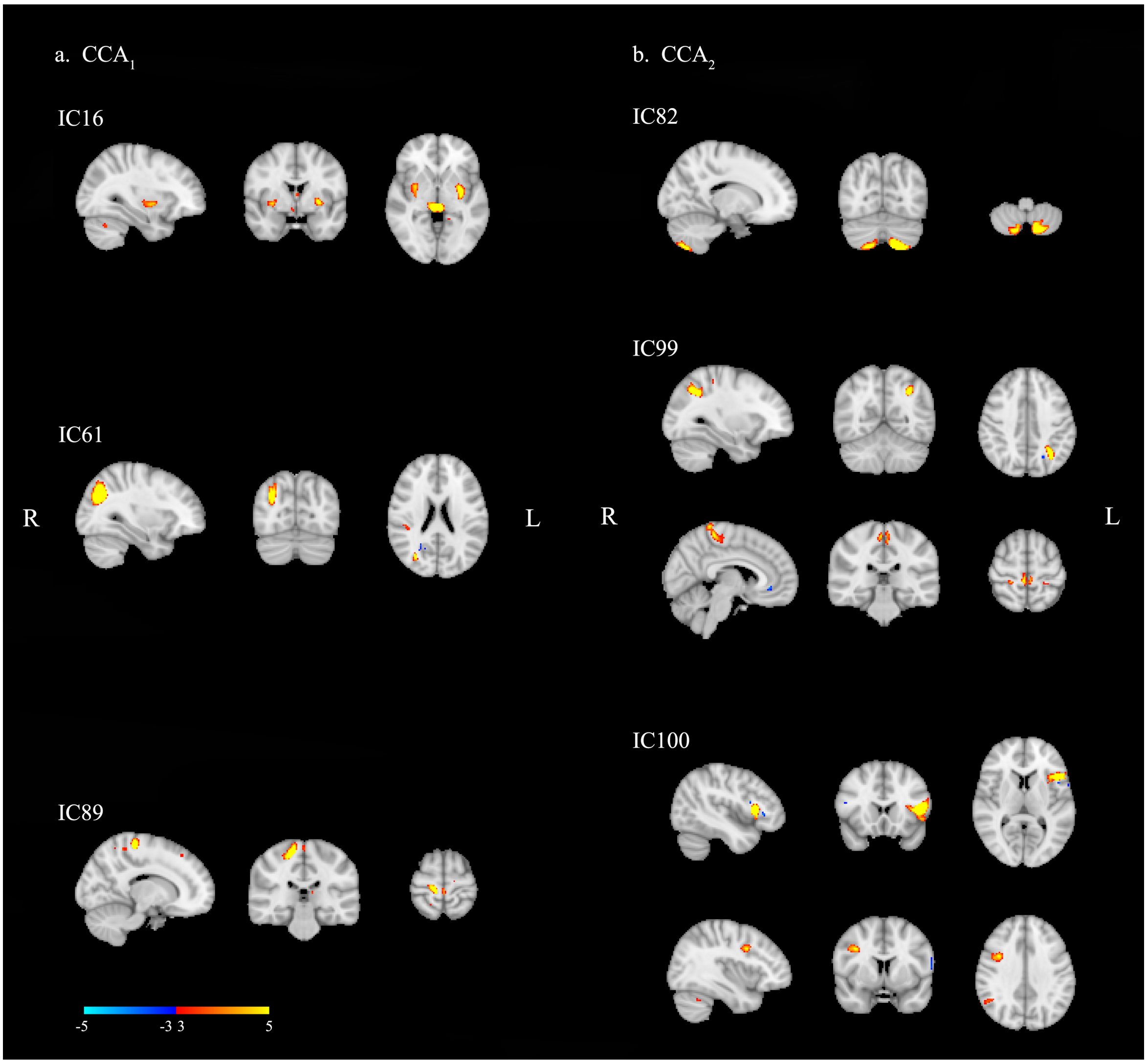

**Figure S6.** Components with highest loadings in CCA. Panel **a** shows the three components with highest loadings in $CCA_1$ (correlation with ADI and ADOS subscales). Panel **b** shows the three components with highest loadings in $CCA_2$ (correlation with SRS, RBS, and SSP). The component maps were thresholded at $3<|Z|<5$.

### 10. Uncorrected main CCA mode loadings of each component

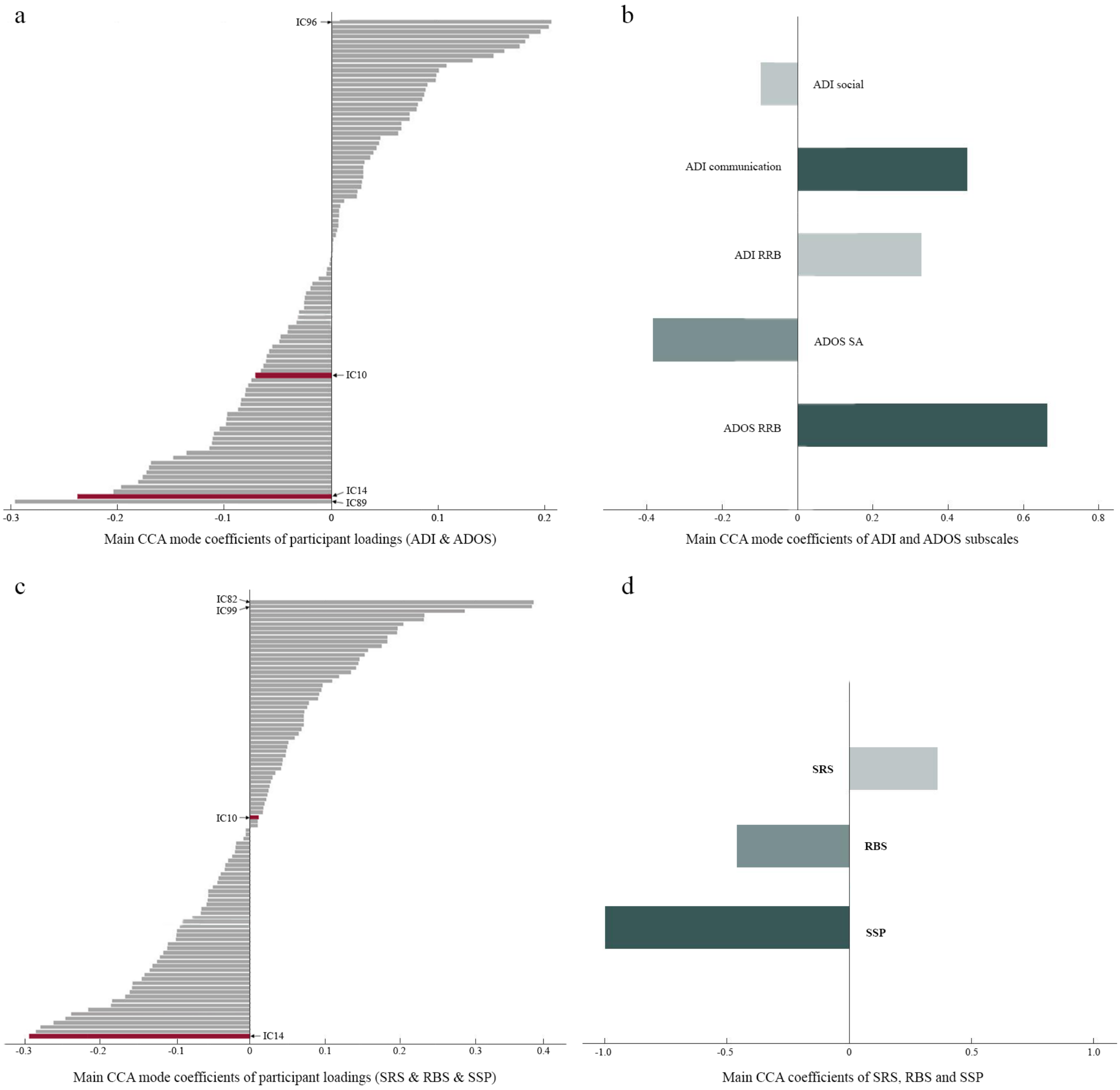

**Figure S7**. Main CCA mode loadings of each component and symptom profiles. (a, c) display the degree that each component contributed to the main CCA mode related with ADI and ADOS (a), and with SRS, RBS, and SSP (c). The two components with significant group effects are displayed in red. (b, d) show the loadings of symptom profiles in each main CCA mode. IC14 ranks third among the 100 components when correlating to ADI and ADOS (a), and it ranks fourth in the CCA with SRS, RBS, and SSP (c). CCA, canonical correlation analysis; ADI, Autism Diagnostic Interview-Revised; ADOS, Autism Diagnostic Observational Schedule 2; SRS, Social Responsiveness Scale 2nd Edition; RBS, Repetitive Behavior Scale-Revised; SSP, Short Sensory Profile; IC, independent component.
